# An endogenous basis for synchronization manners of the circadian rhythm in proliferating *Lemna minor* plants

**DOI:** 10.1101/2021.02.09.430421

**Authors:** Kenya Ueno, Shogo Ito, Tokitaka Oyama

## Abstract

Endogenous circadian rhythms in plants play a role in adaptation to day-night cycles. The circadian clock is a cell-autonomous system that functions through the coordination of time information in the plant body. Synchronization of cellular clocks is based on coordination mechanisms; the synchronization manners in proliferating plants remain unclear. We performed spatiotemporal analysis of the circadian rhythm of fronds (leaf-like plant units) of proliferating *Lemna minor* plants carrying a circadian bioluminescence reporter, *AtCCA1:LUC*. Noninvasive observations of the bioluminescence of fast-growing two-dimensional plants allowed us to analyze the circadian rhythms at a cell-level resolution and obtain information regarding frond lineage. We focused on spontaneous circadian organization under constant light conditions for plants with light/dark treatment (LD-grown) or without it (LL-grown). Even fronds developing from an LL-grown parental frond showed coherent circadian rhythms among them. This allowed the maintenance of circadian rhythmicity in proliferating plants. Inside a frond, a centrifugal phase/period pattern was observed in LD-grown plants, whereas various phase patterns with traveling waves were formed in LL-grown plants. These patterns were model-simulated by local coupling of cellular circadian oscillators with different initial synchronous states in fronds. Taken together with similar patterning previously reported for detached leaves of *Arabidopsis*, it is strongly suggested that local coupling is the primary force for the development of these phase patterns in plants lacking long-distance communication. We propose a basic framework of spontaneous phase patterning with three stages of circadian organization: initial phasing, evolution of patterning, and desynchronization/randomizing of phase, in association with altering cell-cell coupling.

## Introduction

Endogenous circadian rhythms are important for the coordination of plant physiology under environments with 24-h day-night cycles [1,2]. The circadian clock is an endogenous timing system based on self-sustained oscillations in individual cells of plants [3]. The core circadian oscillator is driven as a gene regulatory network comprising clock genes. Many clock genes such as *CCA1, LHY, PRRs, LUX*, and *TOC1* form transcription-translation feedback loops in *Arabidopsis*.

The coordination of time information within the plant has been studied from the aspects of cell-cell local coupling and long-distance communication between tissues [2–4]. With respect to the internal circadian organization of plants, spatial phase patterns have been reported in the leaf (shoot) and in the root under constant conditions [5–11]. It has been suggested that phase attractive interactions exist between neighboring cellular oscillators, although the strength is not sufficient to globally synchronize these organs. Tissue-specific functions for circadian coordination have been reported. Vascular tissues show robust circadian rhythms and have a dominant effect on other tissues [12]. Shoot apexes likely play a dominant role in the regulation of the circadian clock of the whole plant, and the ELF4 gene product mediates shoot-to-root communication [13,14].

During the observation period of circadian rhythms, plants usually grow more or less; however, the circadian organization of the growing plants has rarely been studied. In the elongating roots of *Arabidopsis CCA1:LUC* transformant seedlings, a stripe pattern of bioluminescence can be observed as a traveling wave [8]. It was suggested that the circadian rhythm at the root tip was reset at a developmental stage, and a phase attractive interaction occurred between mature cells. Root elongation occurs one-dimensionally at the root tip; this simple growth makes it easier to study the circadian behavior of the growing organ. In contrast, the growth mechanisms of the shoot are three-dimensionally complex, and it is difficult to experimentally study the circadian behavior.

Duckweed plants proliferate two-dimensionally on the water surface, and their flat and tiny bodies with rapid growth facilitate the monitoring of whole growing plants [4]. The plant body consists of several fronds (leaf-like structures) that form a colony. A frond of duckweed in the *Lemna* genus has two pockets; deep inside these pockets, a meristematic tissue is located [15]. New fronds (daughter fronds) separate from the parental frond during proliferation. By the time separation occurs, daughter fronds are usually matured, and the frond size stops increasing. These developmental features enable us to access the lineage of individual plants (fronds) and positions of individual mature fronds in proliferating plants.

Circadian rhythms in important physiological processes, such as CO_2_ output and potassium uptake, have been well characterized in duckweeds [16–18]. Recently, bioluminescence monitoring systems have been successfully applied to monitor circadian rhythms using a particle bombardment method [19–21]. *AtCCA1:LUC* is a circadian reporter that exhibits a bioluminescence rhythm in duckweed similar to that in the *Arabidopsis* transformant. Luminous cells that are transfected by particle bombardment can be monitored at the single-cell level. Cellular bioluminescence rhythms have been quantitatively analyzed in *Lemna gibba* [9, 22, 23]. In addition to the heterogeneous properties of cellular oscillators, spatial characteristics such as local similarities in phases and a centrifugal phase pattern in fronds have been demonstrated. Because only <100 cells in a mature frond can be transfected by the particle bombardment method [9], this monitoring system is unsuitable for measuring the circadian rhythms of newly emerging fronds and for high-resolution observations.

To reveal the dynamic behavior of circadian rhythms in proliferating plants, we spatiotemporally analyzed the bioluminescence of transgenic *Lemna minor* plants carrying a circadian bioluminescence reporter, *AtCCA1:LUC*. Through comparative analysis between fronds in proliferating plants together with rhythm analysis and mathematical modeling at a resolution close to the cell size, we propose principles of spontaneous circadian organization in plants.

## Results

### Experimental setup

To study the internal circadian organization of proliferating plants, we monitored the bioluminescence of a transgenic *Lemna minor* that carried an *AtCCA1:LUC* reporter gene in the genome (*SI Materials and Methods*). The bioluminescence of proliferating plants was imaged under constant light every hour for more than a week using an automated imaging system with an EM-CCD camera [22,24] (Movies S1-4). We prepared plants grown under 12-h light/12-h dark cycles (LD-grown) or constant light (LL-grown) before monitoring. During bioluminescence monitoring, bright-field images were captured every hour to obtain information on frond contour. Image alignment for individual mature fronds was performed to assign each pixel to a specific position in the frond (*SI Materials and Methods*). Under constant light, the transgenic duckweed plants grew to double the size in 2-2.5 days (Fig. S1*A* and *B*). Newly emerging fronds (daughter fronds) matured in 2 – 3 days and then stopped growing. Daughter fronds also produced new fronds (Fig. S1*C*–*F*). Some daughter and granddaughter fronds moved and became out of sight of the camera during monitoring. Circadian rhythmicity under constant conditions was not observed during the growth of either LD-grown or LL-grown plants (Fig. S1*B*).

### Maintenance of circadian phases of fronds from LD-grown plants under constant light

Bioluminescence of the proliferating plants from an initial colony of LD-grown duckweed was monitored under constant light (Fig. S2*A* and Movie S1). The luminescence intensities (signals pixel^−1^ min^−1^) of each frond were calculated (Fig. 1*A*). On the whole, all fronds showed robust circadian rhythms with similar period lengths and phases (Fig. 1*A* and Fig. S3*A*), Bioluminescence rhythms of fronds of the initial colony had very low levels of luminescence at troughs until 2 – 4 days under constant light, and then, the trough levels gradually increased (Fig. S4*A*). Bioluminescence rhythms of newly emerging fronds also showed very low levels of troughs for the first 4 – 5 days (Fig. S4*B* and *C*), and then, the trough levels gradually increased (Fig. S4*B*). Thus, the offspring of an LD-grown duckweed showed robust rhythms under constant light irrespective of the amplitude of the parental frond rhythm (Fig. S4*D*),

**Fig. 1.**
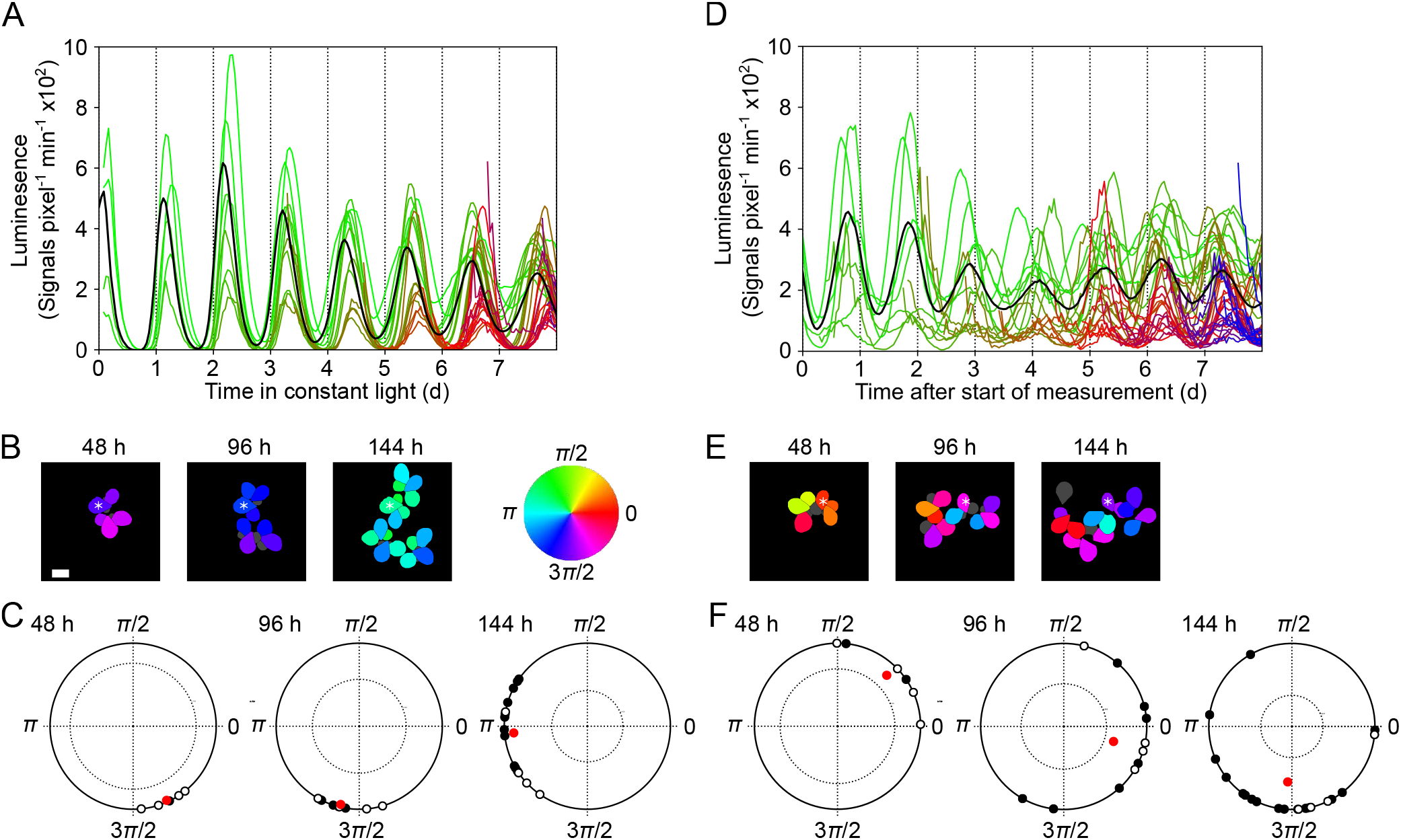
Bioluminescence rhythms and the phase properties of individual fronds. Data of fronds observed in the experiments LDLL1 (A-C) and LLLL1 (D-F) are represented. (A, D) Plots for bioluminescence rhythms of individual fronds under constant light. Color gradients are applied according to the order of emergence of fronds. The black line represents mean values of all fronds. Time 0 denotes the last transition from dark to light (A) or the start of measurement (D). (B, E) Comparison of phases of fronds at 48 h, 96 h, and 144 h. The phases are color-coded. Gray represents fronds the phase of which is not defined. Asterisks represent the oldest frond. Scale bar: 3 mm. (C, F) Plots of phases of fronds on the unit circle at 48 h, 96 h, and 144 h. Open circles on each unit circle represent phases of fronds of the initial colony, and filled circles represent phases of newly emerging fronds. The tip of a vector (red dot) represents the vector mean of each plot on the unit circle. The vector length and the angle represent the SI and a representative phase at each time. A dotted circle inside the unit circle represents a significance level of 95%. The confidence limit was calculated based on random sampling (10,000 repeats).

Phase differences among fronds were analyzed in detail at 48 h, 96 h, and 144 h, when the numbers of measurable fronds were 7, 12, and 21, respectively (Fig. 1*B* and *C*). The maximum phase differences at 48h, 96h, and 144h were 0.57rad [Circadian Time (CT): 2.2h], 0.86rad (CT: 3.3h), and 1.57rad (CT: 6.0h), respectively. Thus, the phase differences gradually increased as plants proliferated from an LD-grown duckweed under constant light. As shown in the unit circle for phase at 144h, circadian rhythms of newly emerging fronds tended to show a phase delay compared with those of fronds of the starting colony (Fig. 1*B* and *C*). This phase delay in newly emerging fronds caused the gradual increase in phase differences. Nevertheless, throughout monitoring, circadian rhythms were highly synchronous among fronds, with a synchronization index (SI) value at 144h of 0.89.

Similar results were obtained for the damping and synchronization phenomena for a different transgenic line under the same experimental conditions (Fig. S3*B*, S5). Thus, the overall synchronization of circadian rhythms between fronds emerging from an LD-grown colony is likely to be maintained under constant conditions.

### Synchrony of circadian rhythms among fronds from LL-grown plants under constant light

The bioluminescence of proliferating plants starting from an LL-grown colony was monitored under constant light (Fig. S2*B*, Movie S3). These plants never experienced entrainment stimuli from the environment; most fronds showed circadian rhythms with lower amplitudes than those of LD-grown plants (Figs. 1*A* and *D* and Figs. S4 and S6). In contrast to LD-grown plants, these fronds with circadian rhythms showed a large difference in peak times. Interestingly, the bioluminescence of the total measured fronds showed a circadian rhythm, suggesting that circadian phases of individual fronds started from an LL-grown colony were not randomly determined (Fig. 1*D*). Phase differences between fronds were analyzed in detail at 48 h, 96 h, and 144 h, when the numbers of measurable fronds were 9, 15, and 21, respectively (Fig. 1*E* and *F*). SI values at 48h, 96h, and 144h were 0.85, 0.63, and 0.67, respectively. These values were significantly larger than those predicted in random phases. This indicated that the circadian phases of individual plants started from an LL-grown colony were coherent to some extent. This also suggested that the circadian phases of newly emerging fronds could be spontaneously determined through their parental colonies to become synchronous with each other.

Similar results were obtained for the synchrony between fronds in a duplicate experiment, which were started with a parental frond showing almost arrhythmic bioluminescence (Fig. S7). This suggests that synchronization between newly emerging fronds might occur irrespective of the circadian states of their parents.

### Centrifugal phase patterning in fronds of proliferating LD-grown plants

As observed in Movie S1, Fig. S2*A*, and *C*, some spatial patterns of bioluminescence rhythms appeared within each frond of LD-grown plants under constant light. To assess spatial patterning, luminescence time series of pixels were analyzed for the circadian phase and period (*SI Materials and Methods*). Fig. S8*A* shows examples of luminescence time series of single pixels in the mature fronds that had ceased to grow. Circadian rhythms with gradual damping were observed in fronds of the initial colony (Fig. S8*A* LDLL1_1, _1-1), This damping appeared to be due to the desynchronization of cellular rhythms in the region around the pixels [9]. Meanwhile, such damping was not observed in a newly emerging frond, whereas its parent showed severe damping (Fig. S8*A* LDLL1_1-1-2). The stability of the cellular circadian rhythms seems to depend on the age of frond.

Circadian phases for individual pixels in the fronds are mapped every 24h (48-168 h) (Fig. 2*A*). As shown in these phase maps, each frond showed a centrifugal pattern in which the phases near the center of the frond were advanced compared with those in the peripheral region. The phase differences were likely caused by a position-dependent difference in the period lengths. Six out of seven fronds showed a high positive correlation between period lengths and the distance from the geometric center of the frond (Fig. 2*A* and *B*, Table S1). Thus, period lengths centrifugally increased by ~1 h in a frond of LD-grown plants. The transition of the phase differences within a frond was quantitatively evaluated for individual fronds with SI *R*(*t*) (Fig. 2*C*). Mature fronds at the start of monitoring, LDLL1_1 and _1-1, exhibited SI values of 0.98 first, which then gradually decreased to 0.74 and 0.41 at 168 h, respectively. Immature fronds at the start (1-2, 1-1-1) and newly emerging fronds (1-3, 1-1-2, and 1-1-1-1) firstly showed an SI of 0.9 or larger, which then gradually decreased. Interestingly, the first SI value of any frond was larger than that of older fronds at that time. This means that a frond always emerged with a higher synchrony than its parent. Consequently, the circadian rhythm of the total fronds of proliferating LD-grown plants is robustly maintained regardless of the desynchronization of individual fronds. It seemed that during the development of a frond from LD-grown plants, circadian rhythms were spatially synchronized and synchrony was maintained. As mentioned previously, the total luminescence of individual fronds showed a high-amplitude rhythm for 4 – 5 days, and then, damping occurred (Fig. S4). This damping of the bioluminescence rhythm was due to the progress of centrifugal phase patterning and local desynchronization between cellular circadian rhythms that appear to be prompted in aged fronds.

**Fig. 2.**
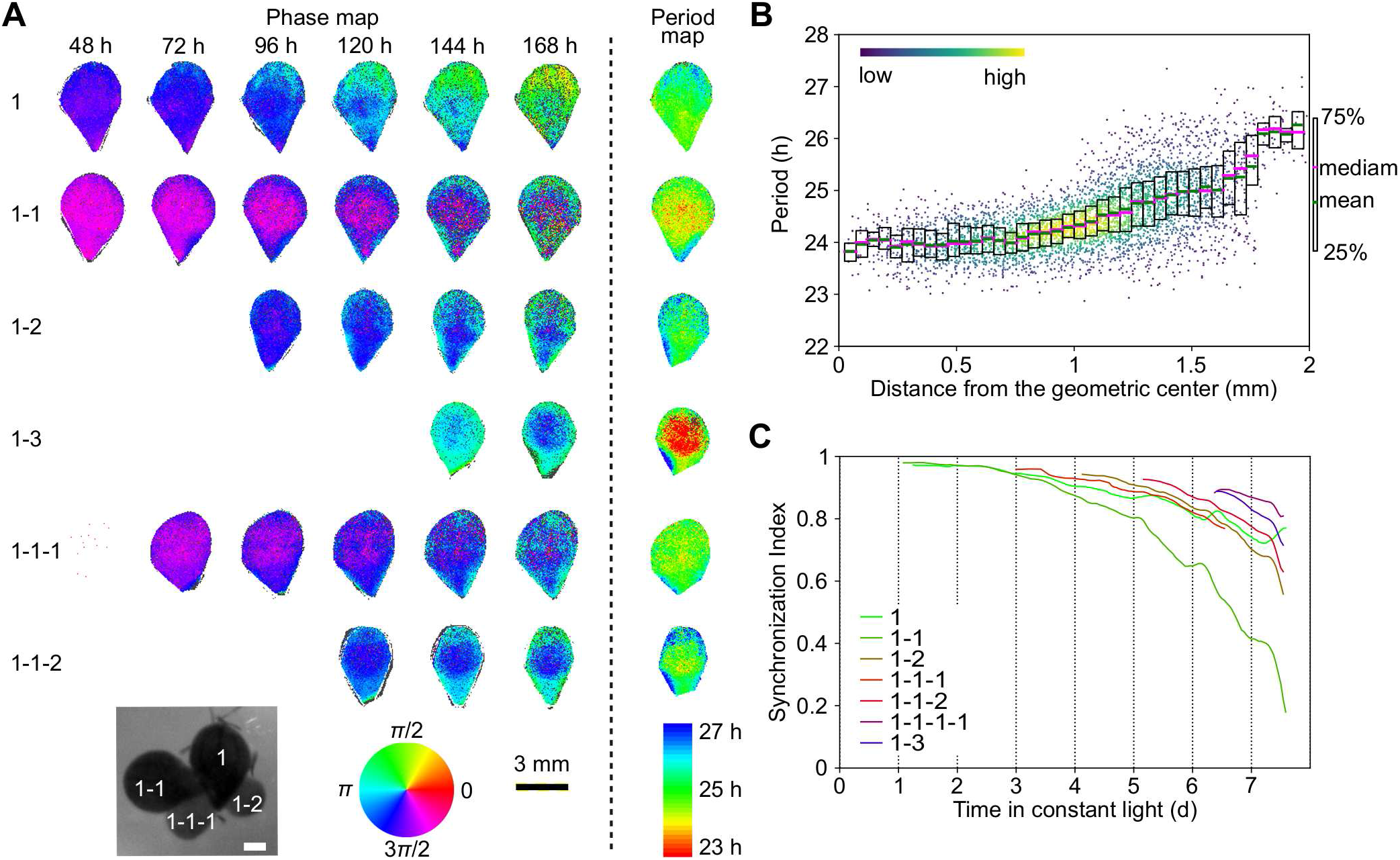
Characterization of bioluminescence rhythms in fronds from an LD-grown plant at the pixel level. The time series of luminescence of individual pixels in the experiment LDLL1 was used for analysis. (A) Phase maps and period maps. Phases at the indicated time and periods of circadian rhythms in each pixel of a frond are color-coded. Frond names are indicated on the left. A frond name has two parts: the number at the end and the remaining part. The number at the end represents the order of emergence from the parent whose name is represented in the remaining part. A top-view image of the initial plant with frond names is shown. The white bar is 1 mm. Period lengths were calculated using time series for 72 h starting from 24 h or the time at which the frond matured. Gray color represents the pixels the phase or period of which is not defined. The scale is indicated by a black bar (3 mm). (B) Relationship between period lengths and distances from the geometric center of the frond (1 – 1) For each pixel, the period length and the distance from the geometric center to its position are plotted as dots. The density of dots is color-coded. Statistics (mean and quartiles) for the period lengths are plotted at every 0.05 mm interval of distances from the geometric center. (C) Temporal changes in SIs of individual fronds.

In an experiment using another transgenic line (LDLL2), the centrifugal phase and period patterns were observed, whereas the degrees of desynchronization of individual fronds were smaller than those of LDLL1 (Fig. S9 and Table S1). This implies that the centrifugal patterning is robust. Indeed, centrifugal patterning was previously reported for fronds of *Lemna gibba* and detached leaves of *Arabidopsis* [6, 7, 9]. Thus, isolated tissues/organs with shapes/sizes similar to fronds of duckweed and *Arabidopsis* leaves may show this pattern in general.

### Various traveling wave patterns of phase formed in fronds of proliferating LL-grown plants

To characterize spontaneously formed spatial patterns of circadian rhythms, we analyzed the luminescence time series of LL-grown fronds at the single-pixel level (Fig. S2*B* and *D*). Fig. S8*B* and *C* shows examples of the luminescence time series of single pixels in mature fronds. All pixels showed circadian rhythms even for fronds showing almost arrhythmic bioluminescence on average (e.g., LLLL2_1). Circadian rhythms with damping were observed in fronds of the initial colony (e.g., LLLL1_1); however, such damping was not obvious in a newly emerging frond (e.g., LLLL1_1-1-2) during monitoring. The stability of circadian rhythms likely depended on the frond age. These features of circadian rhythms in LL-grown fronds at the single-pixel level were the same as those observed in LD-grown fronds, suggesting that the basic features of circadian rhythms in a local region are less affected by the environmental conditions before constant light (Fig. S8).

Despite the similarity in pixel-level circadian rhythms, the spatial patterns of circadian phases in individual fronds were different between LL-grown and LD-grown plants (Figs. 2*A* and 3*A*). As shown in the phase maps for individual fronds, various phase patterns formed by traveling waves were apparent in each frond of LL-grown plants. The following three remarkable traveling wave patterns were observed: centrifugal (LLLL1_1-1), transversal (LLLL1_1-3), and spiral (LLLL1_1-1-1) (Movies S5-7). These spatial patterns were maintained during monitoring, suggesting that each frond had a phase pattern based on its circadian property. These spatial patterns seemed to be stochastically formed during frond development because phase patterns were different between daughter fronds of a single parent (1-1 and 1-3) and between a daughter and the parent (1-1-1 and 1-1). The variation in phase patterns was reflected through various SI values (0.1-0.9) (Fig. 3*B*). The SI values of most fronds showed a decreasing tendency during monitoring. This suggested that cellular circadian rhythms were highly coupled in a young frond, and desynchronization occurred in the mature frond as a whole. As expected from the traveling waves of circadian phases in a frond, even a frond with a low SI value showed high synchrony of local region rhythms in a short range (<0.5mm) (Fig. 3*C*). The linkage between phase differences and spatial distances was also observed in studies at the single-cell level using *L. gibba* [9]. These results indicated a phase-attractive interaction between neighboring cellular clocks.

**Fig. 3.**
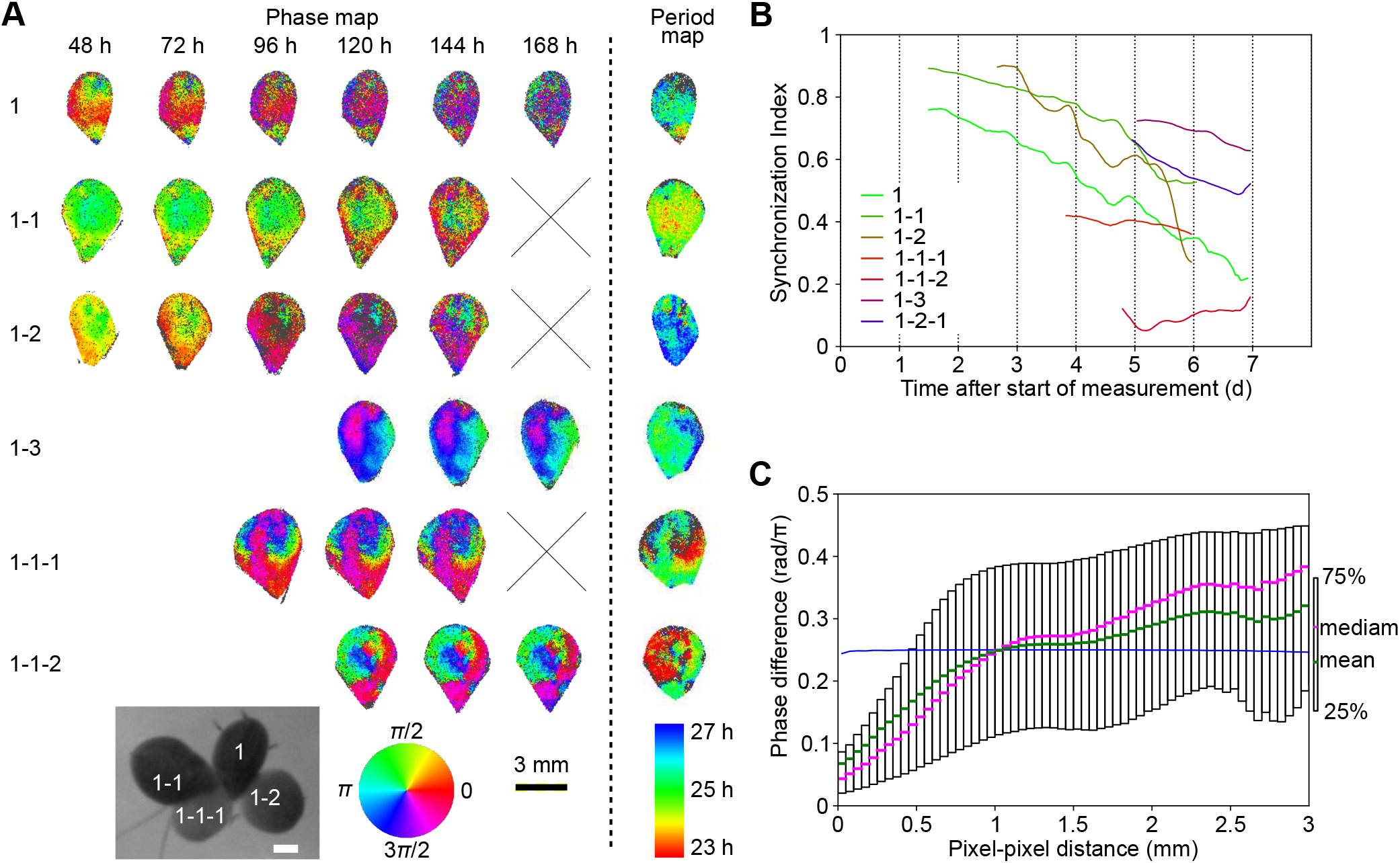
Characterization of bioluminescence rhythms in fronds from an LL-grown plant at the pixel level The time series of luminescence of individual pixels in the experiment LLLL1 was used for analysis. (A) Phase maps and period maps. The legend is the same as Fig 2A. A cross mark represents missing data due to the movement of the frond out of the visual field of the camera. (B) Temporal changes in SI values of individual fronds. (C) Relationship between the phase differences between pixels and pixel-to-pixel distances. Phase differences for all pairs of pixels in the frond (1-1-2) at 120 h and the pixel-to-pixel distances corresponding to the pairs are plotted. Statistics (mean and quartiles) for the phase differences are also shown at every 0.05 mm interval of pixel-to-pixel distances. Blue symbols at each pixel-to-pixel distance interval represent 95% confidence limits of the median, assuming that samples in each interval have the same median value. The confidence limits were calculated based on resampling (10,000 repeats) from all pairs.

Concerning the spatial distribution of the period lengths in a frond, various patterns that seemed to be similar to the phase patterns were detected (Fig. 3*A*). This suggested that the period length at a point was associated with the synchrony in the local area but not with the location per se in the frond. Frond (1-1), which showed the highest synchrony at the start of monitoring, showed a centrifugal phase pattern and shorter periods around the central area than those in the periphery (Fig. 3*A*). The centrifugal patterns of the phase and period were also observed in LD-grown fronds (Fig. 2). This implies that a highly synchronous frond may form a centrifugal pattern irrespective of the light/dark treatment.

Fig. S10 shows the results for a duplicate experiment of LL-grown duckweed (LLLL2). As presented in Fig. 3 for LLLL1, various phase patterns, a large variation in synchrony within fronds (SI values: 0.0-0.8), and a high synchrony in the local region (<0.5mm) were observed.

Taken together with the results from Fig. 1*D-F*, in proliferating LL-grown plants, various phase patterns were observed in individual fronds with high local synchrony before frond maturation, but daughter fronds from a parent shared similar phases on the whole.

### A mathematical model to form phase and period patterns in mature fronds

An interesting feature observed in the spatiotemporal patterns of the bioluminescence rhythm was that LD-grown plants showed a centrifugally increased period pattern in the frond under constant light. This period pattern was accompanied by a centrifugal phase pattern that caused desynchronization in the frond. The centrifugal period pattern was obvious only in fronds with high synchrony at the start of monitoring. Thus, it was unlikely that the natural frequencies of the individual cellular oscillators showed a similar centrifugal pattern. This pattern was likely formed based on cell-cell interactions to modify their frequencies. To simulate the centrifugal phase pattern of oscillators in a one- or two-dimensional region, a coupled oscillator model for modified Poincaré oscillators (twist oscillators) has been proposed [25]. This oscillator exhibited limit cycle behavior with angular velocity modified by its amplitude. This mathematical model was applied to the centrifugal patterning of circadian oscillations in mammalian choroid plexus (CP) tissues in the brain [25]. This model postulated that a phase-attractive interaction occurred between neighboring cellular oscillators. The centrifugal pattern emerged due to differences in the coupling input to the cell between the center and boundary regions. We applied this model of mammalian CP to the simulation for phase/period patterning in duckweed fronds. The simulation was performed for cellular oscillators on a square lattice using a realistic geometry of the frond. As expected, a centrifugal period pattern in the frond was caused by synchronous cellular oscillators in the simulation using a set of parameters (Fig. 4*A* and *B*, and Fig. S11). This result mimics the centrifugal period pattern observed in LD-grown plants (Fig. 2 *A* and *B*, Fig. S9 *A* and *B*). In the simulation starting from asynchronous cellular oscillators with the same set of parameters, a phase pattern with spiral traveling waves and a period pattern associated with the phase pattern was observed for the frond (Fig. 4*A* and Fig. S11). The phase/period patterns were observed in some fronds of LL-grown plants (LLLL1_1-3, _1-1-1, _1-1-2 in Fig. 3*A*). Furthermore, the relationship between the phase differences between pixels and pixel-to-pixel distances was quantitatively simulated (Figs. 3*C* and 4*C* and Fig. S10*C*).

**Fig. 4.**
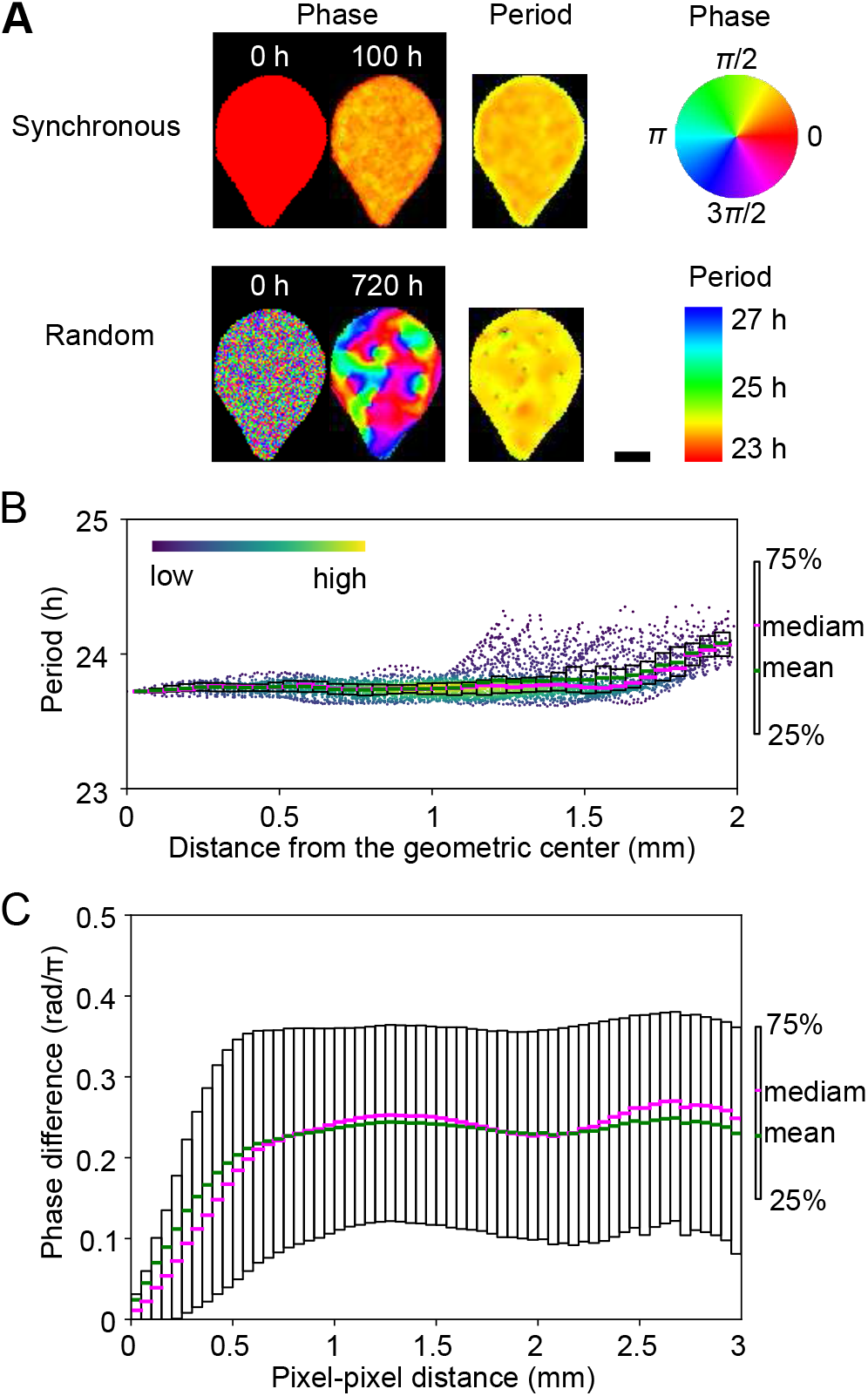
A model simulation for phase and period patterns in fronds. A two-dimensional lattice of nearest-neighbor coupled oscillators on a realistic geometry of a frond is analyzed. A total of *N* = 4809 grid cells (cell size = 40 μm) were placed with respect to the geometry. (A) Phase- and period maps for a frond. Initial phase patterns for synchronous- and randomly phased oscillators were set for simulation with the same coupling parameters. Phases at the indicated time and periods of circadian rhythms in each pixel of a frond are color-coded. Scale bar: 1 mm. (B) Relationship between period lengths and distances from the geometric center of the “synchronous” frond at 100 h. For each pixel, the period length and the distance from the geometric center to its position are plotted as dots. The density of dots is color-coded. Statistics (mean and quartiles) for the period lengths are plotted at every 0.05 mm interval of distances from the geometric center. (C) Relationship between the phase differences between pixels and pixel-to-pixel distances. Phase differences for all pairs of pixels in the “random” fronds at 720 h and pixel-to-pixel distances corresponding to the pairs were calculated. Statistics (mean and quartiles) for the phase differences are plotted at every 0.05 mm interval of pixel-to-pixel distances.

Thus, the coupled oscillator model fits phase and period patterns in both LD-grown and LL-grown plants. This suggests that the phase-attractive interaction between neighboring cells is the driving force for spontaneously organizing cellular oscillation behavior in the frond under constant light.

## Discussion

By monitoring *L. minor* transformants expressing circadian bioluminescence, we performed spatiotemporal analysis of spontaneous circadian organization of proliferating plants under constant light. Spatial patterning of circadian rhythms in a frond and the relationship of circadian phases between fronds from a colony were observed from the data of two pretreatment conditions (LD-grown and LL-grown). With respect to the circadian organization in the duckweed, we propose the following two rules. Local cell-cell interactions are largely responsible for the spatial patterning in fronds. Time information is transmitted from a parental frond to daughters and is maintained in proliferating plants. These rules are applied to duckweed plants with various synchronous states, irrespective of the pretreatment conditions.

Cell-cell interactions for local and spontaneous synchronization of circadian rhythms in plant tissues have been proposed previously [6–10]. Local coupling is presumed to contribute to phase patterns observed in the leaves and roots of *Arabidopsis* transformants expressing circadian bioluminescence. From our previous study of single-cell bioluminescence rhythms in a duckweed, *L. gibba*, weak cellcell coupling was proposed to be associated with the high synchrony of local region rhythms in a short range [9]. By observing transgenic *L. minor* at a high spatial resolution, a centrifugal pattern of circadian rhythm in fronds of LD-grown plants under constant light was demonstrated. The phase pattern was also referred to in an *L. gibba* frond under light/dark cycles [9]. Interestingly, the centrifugal pattern was reported for detached leaves from LD-grown *Arabidopsis* under constant light [6, 7]. In *Arabidopsis*, detached leaves tended to show different circadian patterns from those of intact plants, suggesting long-range transmission of time information to a leaf through vascular tissues. Because vascular tissues are degenerated in duckweed plants, many tissues in mature fronds appear to be isolated from other parts of the colony. This suggests that circadian patterns found in the frond are formed mainly through local cell-cell interactions, as proposed for the circadian patterns in *Arabidopsis* detached leaves. In a population of oscillators with local coupling, spatial phase patterning depends on the network structures, which include the boundary condition and population size [26]. Because the shape and size are relatively similar between duckweed fronds and the observed *Arabidopsis* leaves, centrifugal patterns in these organs may be formed in a similar manner to local coupling.

To interpret the centrifugal pattern in fronds, we applied a coupled oscillator model for a population of modified Poincaré oscillators (twist oscillators) [25]. This model was successfully applied to the centrifugal phase patterning of circadian oscillation in the mammalian choroid plexus (CP). The model with a set of parameters successfully simulated not only the centrifugal phase/period patterns of LD-grown plants but also the spiral traveling wave patterns of LL-grown plants (Fig. 4). In mammalian CP, gap junctions mediate the local coupling between neighboring cells [25]. In plants, plasmodesmata may be responsible for the mediation as gap junctions. Because circadian mediators that transmit the plasmodesmata, ions such as Ca^2+^, low molecular weight metabolites, and clock-related proteins such as *ELF4* may have functions in local coupling [14, 27–29]. The amplitude of each oscillator is an element that modulates its angular velocity (and accordingly, its period length) in this model. In our model, we hypothesized that the synchronous state of the neighboring cells is the amplitude. Higher synchronous states result in a higher frequency (shorter period) than the natural frequency of the oscillator. The centrifugal patterns are unlikely in the coupled oscillator models for amplitude-less phase oscillators (Kuramoto model) and oscillators with periods independent of amplitude (untwisted oscillator) [25]. In a synchronous frond, the central region shows shorter periods than the peripheral regions because cellular oscillators along the boundary of a frond have smaller numbers of neighboring cells than those inside the frond. Following this model, the centrifugal pattern spontaneously appears under constant conditions in fronds with highly synchronous states irrespective of how the way in which synchronization has occurred. In fact, a frond of an LL-grown plant showed a centrifugal pattern (Fig. 3). In contrast, in an asynchronous frond, periods of oscillators are determined depending on the synchrony of neighboring cells without the boundary effect; periods of oscillators largely depend on the spatial phase patterns (Fig. 4*A*). Such phase-pattern-dependent period determination may cause the “after effect” of entrainment on the period length under constant conditions [30]. In *Arabidopsis*, it was reported that photic and temperature entrainment resulted in a difference in periods and phase relation between two circadian outputs [31]. Synchronous states in the regions that express the outputs might become different after entrainment.

In our study, spontaneous circadian organization in proliferating plants was clearly observed; newly emerging fronds of a colony tended to maintain similar phases between them not only in LD-grown plants but also in LL-grown plants. This circadian organization seems to contrast with that observed in *Arabidopsis* growing roots [8]. During development at the root tip, each cell seems to reset its circadian clock at an elongation stage; a traveling wave with a stripe phase pattern is formed along the root. Meanwhile, various spatial patterns were formed in newly emerging fronds from LL-grown plants, suggesting that local coupling between cellular oscillators was important for their phase determination. The coupling between young frond primordia and their parental fronds is presumed to contribute to the overall synchrony among daughter fronds. Various spatial patterns in newly emerging fronds of LL-grown plants appear to be formed during developmental processes in their parents with low spatial synchrony. The high spatial synchrony of LD-grown fronds reflects the uniform phase pattern of newly emerging fronds. The maintenance of high synchrony of newly emerging fronds strongly suggests a high synchronous state of frond primordia at an early developmental stage. The shoot apex of *Arabidopsis* seedlings showed a more robust circadian rhythm than other organs [13]. The meristematic tissue for fronds may also show a robust circadian rhythm that results in the maintenance of high synchrony in LD-grown duckweed. The robustness seems to be due to coupling between cells in individual meristematic tissues in the parent. However, it is unlikely that the coupling is so strong as to spontaneously synchronize the frond primordia because newly emerging fronds of LL-grown plants showed relatively low synchrony within the frond.

Taken together, we propose a framework for spontaneous phase patterning in the fronds of proliferating plants (Fig. S12). During the life of a frond, three successive stages for pattering occur: initial phase patterning, evolution of patterning, and then, desynchronization/randomizing of phases. In early development, a phase pattern is formed by parent-daughter interactions and cell-cell coupling in the frond primordium. After emerging from the parent, phase patterning occurs based on the initial phase pattern for more than a week. This patterning is driven mainly by cell-cell coupling. Desynchronization and randomization of phases occurs with aging. Cell – cell coupling may decrease in an old frond, which contributes to the disorganization of phases in the frond. Thus, we propose that the dynamics of the circadian organization in fronds is led based on local cell-cell coupling.

Unlike duckweed, most flowering plants including *Arabidopsis* fully develop vascular tissues. It has been suggested that these tissues are strongly involved in the circadian organization of the plant body through long-distance signal transduction of time information and their dominant role in the pacing of the rhythm in leaves [6, 12, 14]. Thus, the circadian organization of duckweed plants is regarded as a simplified model of flowering plants. Local coupling of circadian rhythms observed in proliferating duckweed is likely associated with the internal circadian organization of growing plants in general. Based on local coupling, circadian organization in individual plants should dynamically occur by their structural characteristics.

In this study, we clearly demonstrated that circadian phases and periods of individual fronds under constant conditions were affected by their developmental stages and synchronous states. Circadian rhythms of gene expression at the whole plant level could be affected by the tissue-organ specificity as well as the developmental conditions of measured plants. In the case of duckweed, the circadian phases of newly emerging fronds are delayed compared to those of their parent (Fig. 1*A-C*). This could be a factor contributing to the broadening of the peak and trough of the rhythm of proliferating plants. As developmental-stage-specific genes with circadian rhythmicity, *chlorophyll a/b binding protein* genes (*CAB* genes) tend to be highly expressed in young growing leaves, and at least some of them showed a circadian gene expression [32]. Thus, circadian rhythms of these genes in proliferating plants may show longer period lengths than those without developmental stage specificity. Furthermore, treatment of growth regulatory compounds, such as phytohormones, could affect circadian properties at the whole plant level indirectly by altering the development of the plant. In fact, it was reported that multiple phytohormones influenced parameters of the plant circadian clock [33]. Examining various circadian reporters with developmental stages or tissue specificity should clarify the circadian organization in the proliferating plant. As a plant material for experimental and theoretical approaches, duckweed plants expressing bioluminescence reporters are promising because they can be easily monitored through spatiotemporal imaging at various organizational hierarchies—single cells, tissues, and plant bodies—in the proliferating plant [9, 22].

## Methods and Materials

### Plant materials and growth conditions

Transgenic *Lemna minor* (strain 5512) lines were maintained in NF medium with 1% sucrose under constant light conditions in a temperature-controlled room (25 °C), as previously described [21]. White light (30 μE/m^2^s) was supplied by fluorescent lamps (FLR40SEX-W/M/36-HG, NEC). Plants were grown on 60 mL of the medium in 200-ml Erlenmeyer flasks plugged with cotton. An incubator (MIR-153, Panasonic Healthcare) was used for the light-dark culture. In this incubator, white light 30 μE/m^2^s was supplied using fluorescent lamps (FL20SS-W, Mitsubishi Electric), and the growth temperature was maintained at 25 °C. A single colony in the culture was used for each experiment.

### Production of transgenic lines

To generate a transgenic *Lemna* plant that possesses a firefly luciferase bioluminescence reporter gene driven by an *Arabidopsis CCA1* promoter, a *pR4GWB501* binary transformation vector harboring a Hygromycin-resistant selectable marker [34] was employed. First, a luciferase (LUC+ Promega) coding sequence split by an intron sequence, *intronLUC+* [21], was cloned into a *pENTR/D-TOPO* (Invitrogen) vector. Then, the *AtCCA1ex4* promoter [35] sequence was cloned into a *pENTR5’-TOPO* (Invitrogen) vector. Finally, a gateway-based tripartite LR reaction was performed with two entry vector plasmids, possessing the *AtCCA1ex4* promoter and the *intronLUC+* coding region, and the binary vector *pR4GWB501* to form the *pAtCCA1ex4:intronLUC+* expression cassette using Gateway LR Clonase II (Invitrogen). The corresponding construct, *pR4GWB501 pAtCCA1ex4:intronLUC+*, was transformed into the *Agrobacterium tumefaciens* (EHA105 strain). Transformation of *Lemna minor* was performed according to Chhabra et al. [36] with some modifications. Calli were induced from *Lemna minor* on 1x Murashige and Skoog agar medium containing 3% sucrose (MS3S) supplemented with 50.0 μM 2,4-Dichlorophenoxyacetic acid (2,4-D) and 5.0 μM Thidiazuron (TDZ). Then, they were co-cultured with Agrobacterium harboring the binary vector on 1x MS agar medium (1% sucrose) supplemented with 1.0 μM 2,4-D, 2.0μM 6-Benzylaminopurine (BAP), and 200 μM acetosyringone for 3 days. The calli were washed several times, and the transformed cells were selected on 1x Gamborg’s B5 agar medium containing 1% sucrose supplemented with 10mg/L hygromycin B 200mg/L cefotaxime sodium salt, 25 μM indole-3-acetic acid (IAA), and 5 μM kinetin; the calli regenerated into fronds. Several independent *Lemna* transformants with strong LUC+ activity were obtained. We chose #1 and #3 transgenic lines that showed no differences in morphology from its parental *Lemna minor* 5512 plants. The transgenic line #3 was only used in the experiment of LDLL2.

### Experimental setup and bioluminescence monitoring

Methods for bioluminescence imaging of *L. minor* transgenic plants were the same as those for single-cell bioluminescence imaging of duckweed as described previously [9, 37]. A single colony of *L. minor* was transferred to a 35-mm polystyrene dish with 4 mL growth medium (NF with 1% sucrose) containing 0.1 mM D-luciferin. For imaging, the parental frond of the colony was anchored with several pins (Austerlitz Insect Pins, stainless steel 0.1mm diameter; Entomoravia, http://entomoravia.eu/). An EM-CCD camera (ImagEM C9100-13; Hamamatsu Photonics, http://www.hamamatsu.com/) was used to detect luminescence. In experiments using LDLL1, LLLL1, and LLLL2, a macro zoom microscope (MVX-10 with an MVPLAPO 0.63 X lens; Olympus Optical) with the EM-CCD camera system was used [37]. In the LDLL2 experiment, a camera lens (XENON 0.95/25MM C-mount; Schneider Optics, http://www.schneideroptics.com/) with the EM-CCD camera system was used [9]. Two bioluminescence images were captured every 60 min with a 240 s exposure. Some bioluminescence images contained spike signals caused by cosmic rays. To remove cosmic ray spikes, the minimum value for each pixel among the two sequential images at a time was used for analysis. After the capturing the bioluminescence, a bright field image was captured for extraction of the frond area.

### Extraction of contour of each frond in a colony

Bright field images were used to extract the contour of each frond in a colony. The contours were roughly determined by combining the following image processing methods: adaptive thresholding for the binarization of frond area, edge detection by the Canny edge detector, and the Sobel filter. Image processing was performed using ImageJ or OpenCV using Python3. Incorrect/lacking contours were manually corrected using ImageJ. Instead of the image processing methods, we also used pix2pix [38] to extract the contours in Python3/Tensorflow. Image pairs of bright field images and manually determined frond areas were used for training. The output for the frond area of an input image was manually corrected in ImageJ. The geometric center and distances from the center were calculated using Python3.

### Acquisition of luminescence intensities and areas of frond

Tracking of each frond in the timeseries images was performed using the contour data. Frond areas and integrated luminescence intensities for a frond were calculated. Computation processing was performed on Python3.

### Acquisition of timeseries data of luminescence intensities of each pixel

The position of a mature frond terminating the growth expansion was corrected using its contours in the time series images. Position correction was performed using findTransformECC [39] in Python3. The time series data of luminescence intensities for each corresponding pixel were acquired from a series of bioluminescence images. For data analysis, we used pixels that had 48 h or more successive data. This excluded pixels near the contour of the frond.

### Data analysis

For data analysis of luminescence intensities of both fronds and pixels, the same methods as described in our previous paper [9] were used to estimate the phase and period.

Peaks in each bioluminescence rhythm were detected as follows. First, time series data were smoothed with the 3-h moving average, and the peak positions were roughly identified as local maxima. The precise positions of the peaks were estimated via local quadratic curve fitting for the smoothed time series. The width of the local fitting area was set to 5 h.

We introduced phase *θ*(*t*) of the bioluminescence rhythms *x*(*t*) as follows

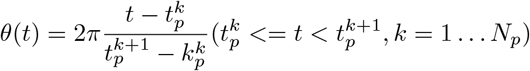

where 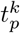 represents the occurrence time of the *k*th peak of *x*(*t*) and *N_p_* represents the total number of peaks.

The period of each luminescence rhythm was estimated by FFT-NLLS, a multicomponent cosine fit [40]. In FFT-NLLS, the rhythm significance was estimated using an RAE that increases from 0 to 1 as the rhythm nears statistical insignificance. Rhythms with a period of 16 – 32 h were taken similarly to those in the circadian range. The FFT-NLLS algorithm was implemented as Python3 scripts [9, 40]. With respect to the periods of pixel luminescence, only data with RAE values less than 0.3 were used.

### Synchronization Index

The synchronization index *R*(*t*), which is known as the order parameter in the Kuramoto model, was calculated at each time point as follows:

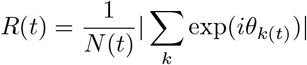

where *θ*_*k*(*t*)_ is the phase of the *k*th frond (or pixel) at time *t*, and *N*(*t*) is the number of fronds (or pixels) [9, 26].

### Mathematical modeling

We adopted a coupled oscillator model (2D model) described by Myung et al. [25].The following equations describe the modified Poincaré oscillator with a coupling factor:

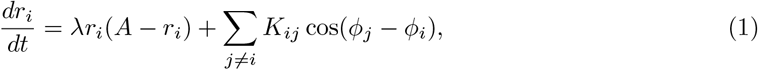

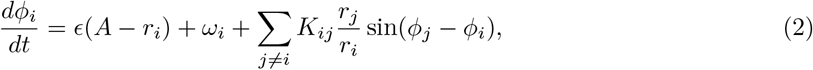

where λ, *A, ϵ, ω_i_*, and *τ_i_* are the radial relaxation rate, amplitude, twist factor, angular velocity (2*π*/*τ_i_*), and the internal free-running period of oscillator *i. K_ij_* is the coupling constant of oscillator *i* to oscillator *j*. Eq. 1 and 2 describe the dynamic evolution of the radial component *r_i_*(*t*) and phase component *ϕ_i_*(*t*). Parameters used for the simulations were *K* = 0.01, *ϵ* = −0.02, λ = 0.05. Identical simulations were run for random sets of Gaussian period (*μ* = 27.0*h*, *σ* = 0.5*h*) and initial phases were set to 0 or were randomized. The interaction for a cell was given by a Moore neighborhood, that is, the eight cells surrounding the cell. Period lengths at a time were represented as the peak-to-peak interval, including the time point.

## Data availability

Python3 analysis codes are available at https://github.com/ukikusa/circadian_analysis/releases/tag/v0.9, https://github.com/ukikusa/twist_simulater, https://github.com/ukikusa/pix2pix_tensorflow. Further data are available on request from the corresponding authors.

## Acknowledgments

We thank Dr. Tsuyoshi Nakagawa for providing us with pR4GWB501 vectors. We thank Dr. Masaaki Morikawa for providing us with *Lemna minor 5512*. This work was supported in part by the Japan Society for the Promotion of Science KAKENHI (Grant numbers 25650098, 17KT0022, and 19H03245), the Japan Science and Technology Agency (JST) ALCA to, and KAKENHI (Grant number 20K06342) to SI. We also thank the MACS program (Graduate School of Science, Kyoto University) for giving us the opportunity for discussions. We would like to thank Editage (www.editage.com) for English language editing.

## Author contributions

Author contributions: K.U., S.I. and T.O. designed the research; K.U. and S.I. performed the research; K.U. analyzed the data; and K.U., S.I., and T.O. wrote the paper.

## Supplementary Information

**Fig. S1:**
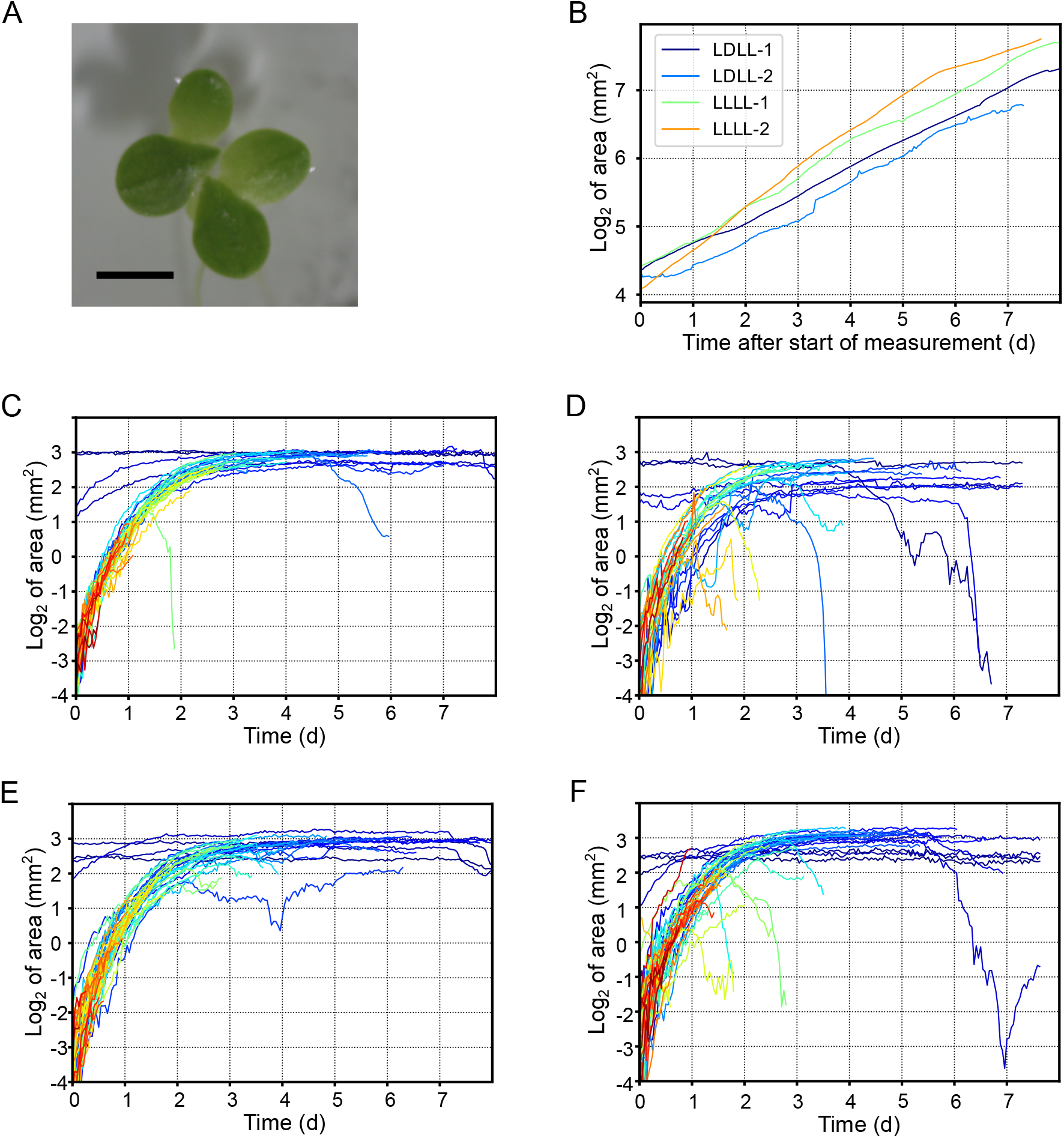
Growth of duckweed plants during the bioluminescence monitoring experiments: LDLL1, LDLL2, LLLL1, and LLLL2. (A) Top view of a colony of the *L. minor* transformant *AtCCA1::LUC#1*. The scale is indicated by a black bar (3 mm). (B) Growth curves of duckweed plants represented by surface area. (C – F) Growth curves of each frond in LDLL1 (C), LDLL2 (D), LLLL1 (E), and LLLL2 (F). Color gradients are applied according to the order of emergence of fronds.

**Fig. S2:**
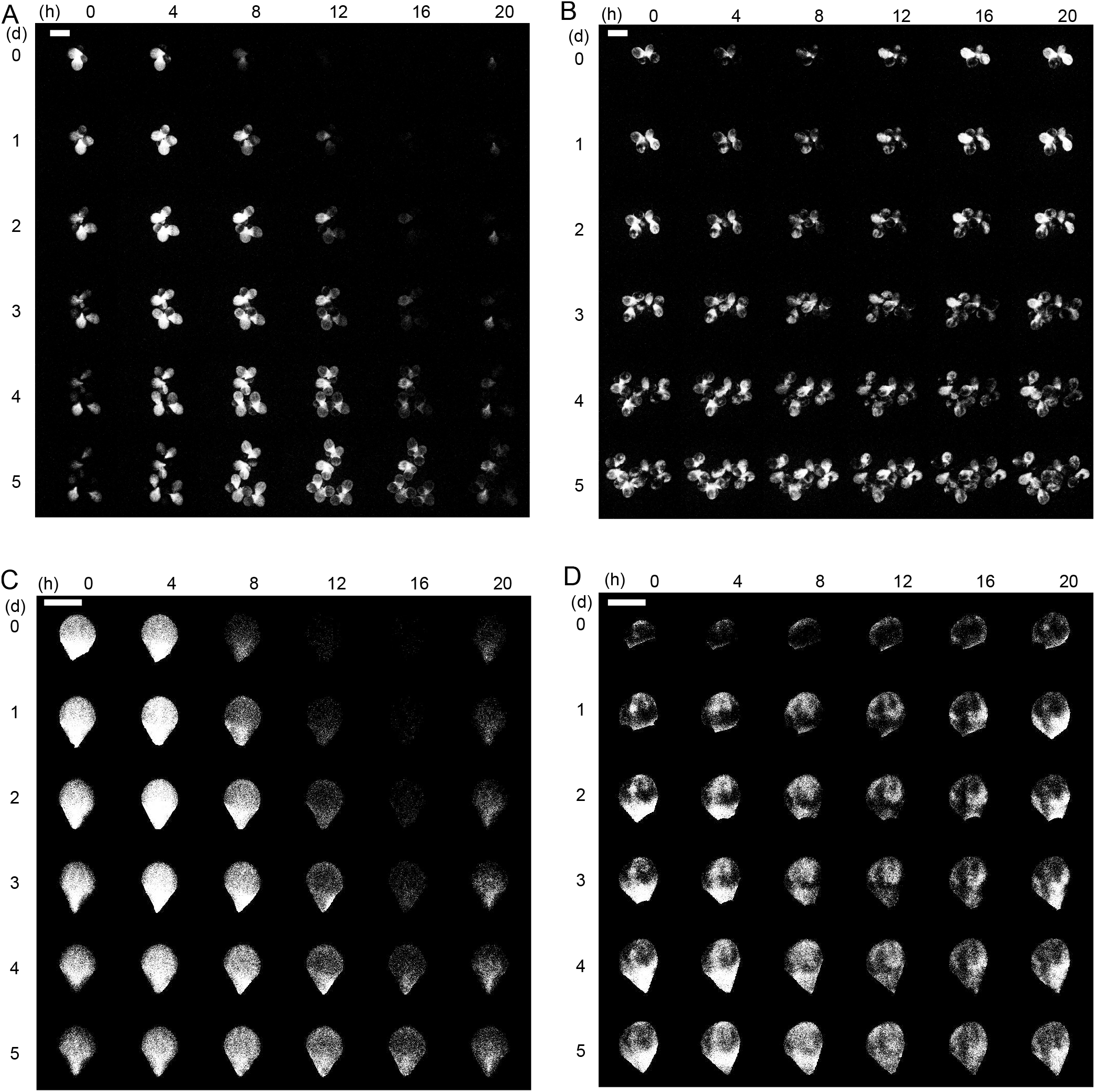
Time-series bioluminescence images of *L. minor* transformant *AtCCA1:LUC*. Images of proliferating plants [(A) LDLL1 and (B) LLLL1] and of a frond [(C) LDLLF_1-1 and (D) LLLLF_1-1-1] every 4 h are sorted for 6 days. Scale bars: 5 mm in (A) and (B) and 3 mm in (C) and (D).

**Fig. S3:**
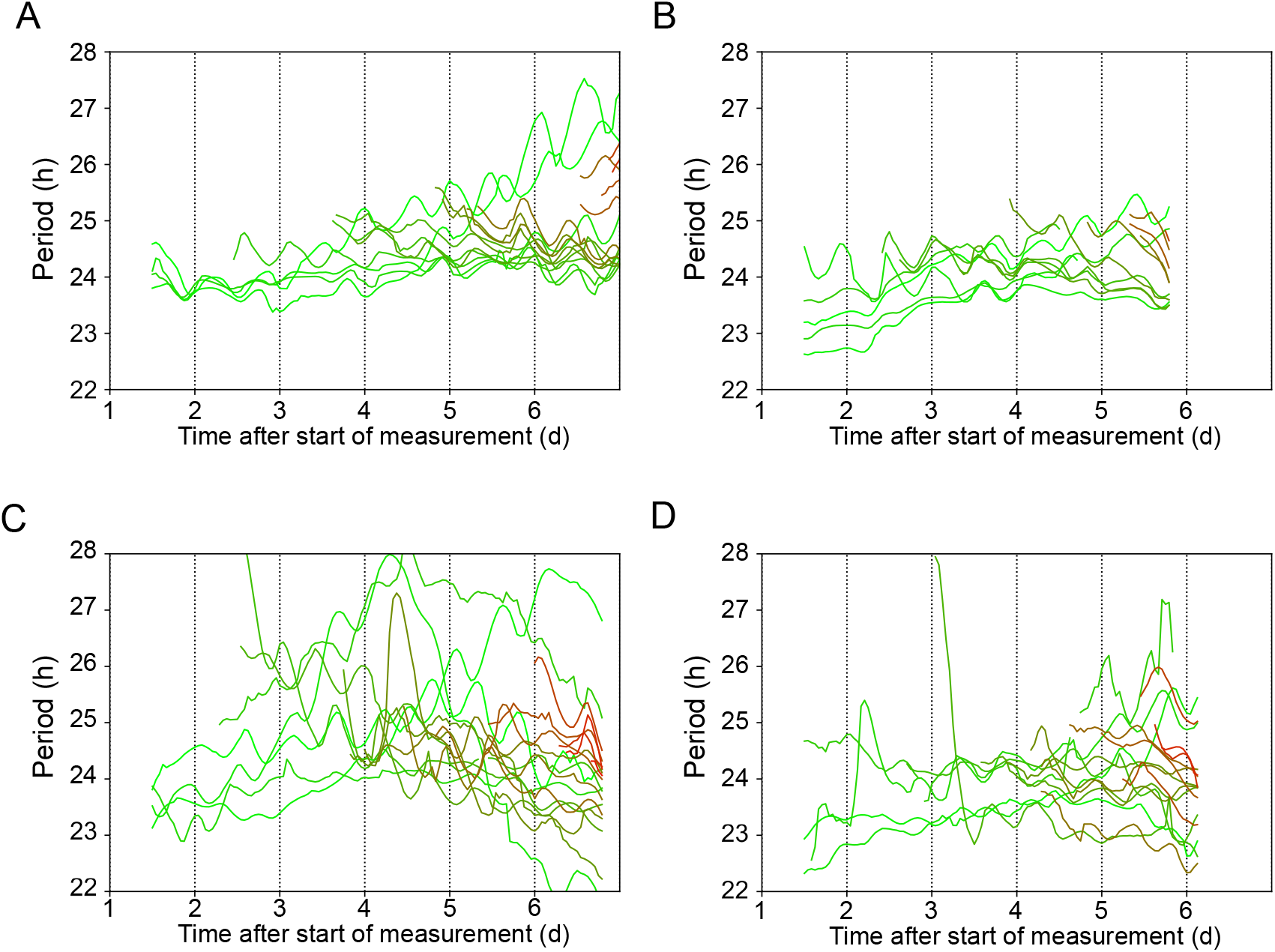
Drifting of period lengths of individual fronds. The period at each point was calculated using FFT-NLLS in the 72 h range (36 h before and after the time). (A) LDLL1, (B) LDLL2, (C) LLLL1, and (D) LLLL2.

**Fig. S4:**
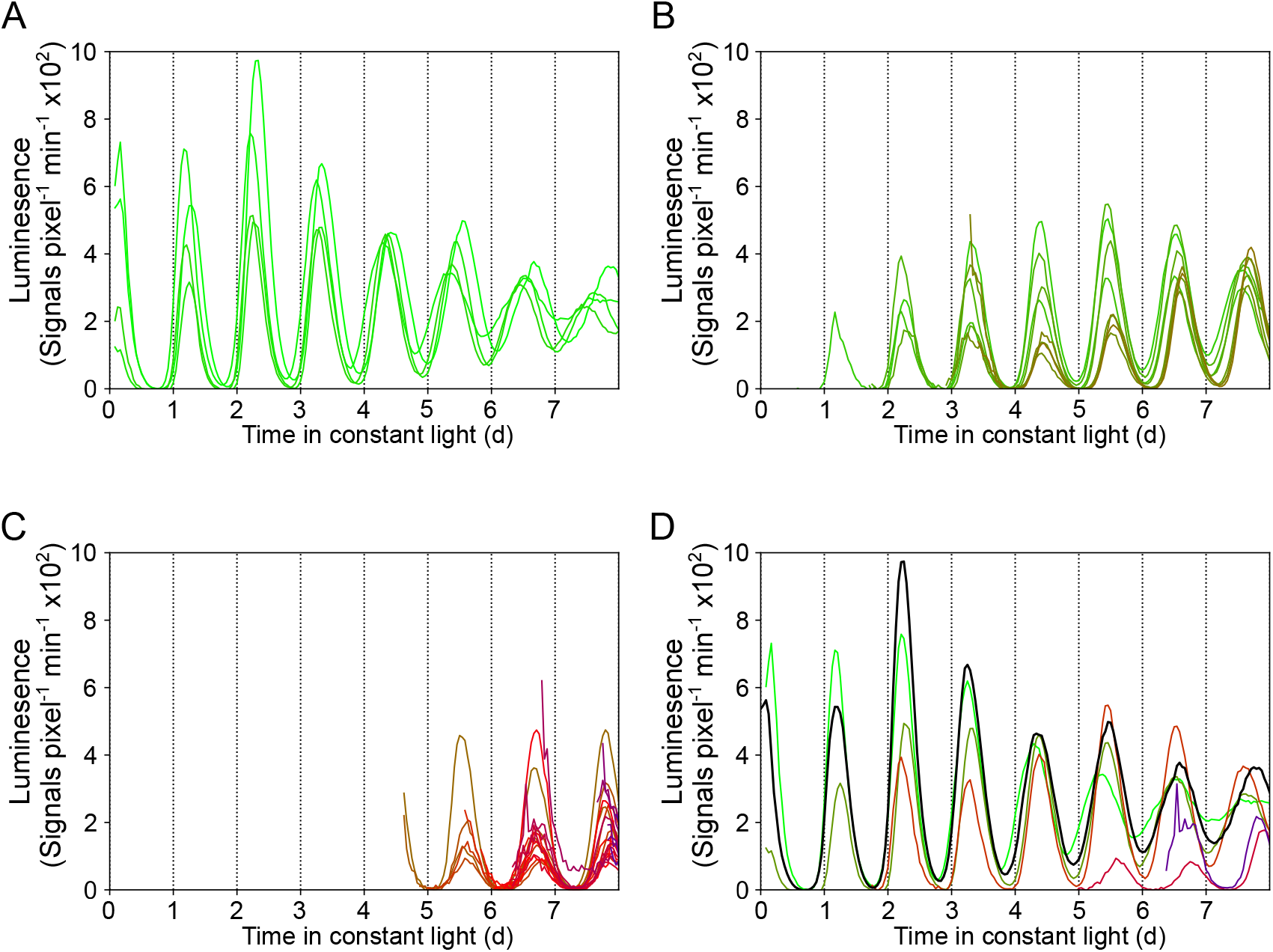
Bioluminescence rhythms of individual fronds of similar ages from an LD-grown plant in the experiment LDLL1. The bioluminescence rhythms plotted in Fig. 1A are classified into those of (A) fronds of the initial colony, (B) newly emerging fronds in the first 4 days, and (C) newly emerging fronds after 4 days. (D) Bioluminescence rhythms of a parental frond and its daughter fronds. The black and colored lines represent the rhythm of the parent, LDLL1_1, and those of its daughters (1-1, 1-2, 1-3, 1-4, 1-5), respectively. Color codes for (A-C) are the same as those in Fig. 1A. Colors in (D) are recoded.

**Fig. S5:**
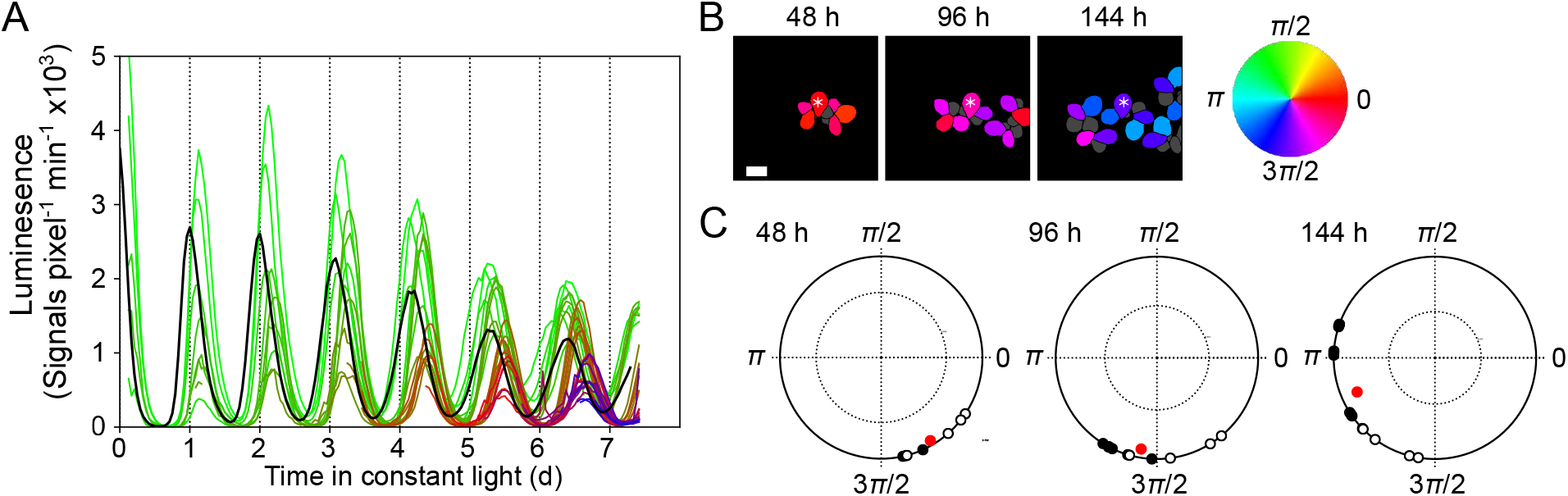
Bioluminescence rhythms and their phase properties of individual fronds from an LD-grown plant in the experiment LDLL2. (A) A plot of bioluminescence of individual fronds under constant light. Color gradients are applied according to the order of emergence of fronds. The black line represents the mean values of all fronds. Time 0 is defined as the last transition from dark to light. (B) Comparison of phases of fronds at 48 h, 96 h, and 144 h. The phase is color-coded. Gray represents the fronds the phase of which was not defined. Asterisks represent the oldest frond. Scale bar: 3 mm. (C) Plots of phases of fronds on the unit circle at 48 h, 96 h, and 144 h. Open circles on each unit circle represent phases of fronds of the initial colony, and filled circles represent phases of newly emerging fronds. The tip of a vector (red dot) represents the vector mean of each plot on the unit circle. The vector length and angle represent the SI and the representative phase at each time, respectively. A dotted circle inside the unit circle represents a significance level of 95%. The confidence limit was calculated based on random sampling (10,000 repeats).

**Fig. S6:**
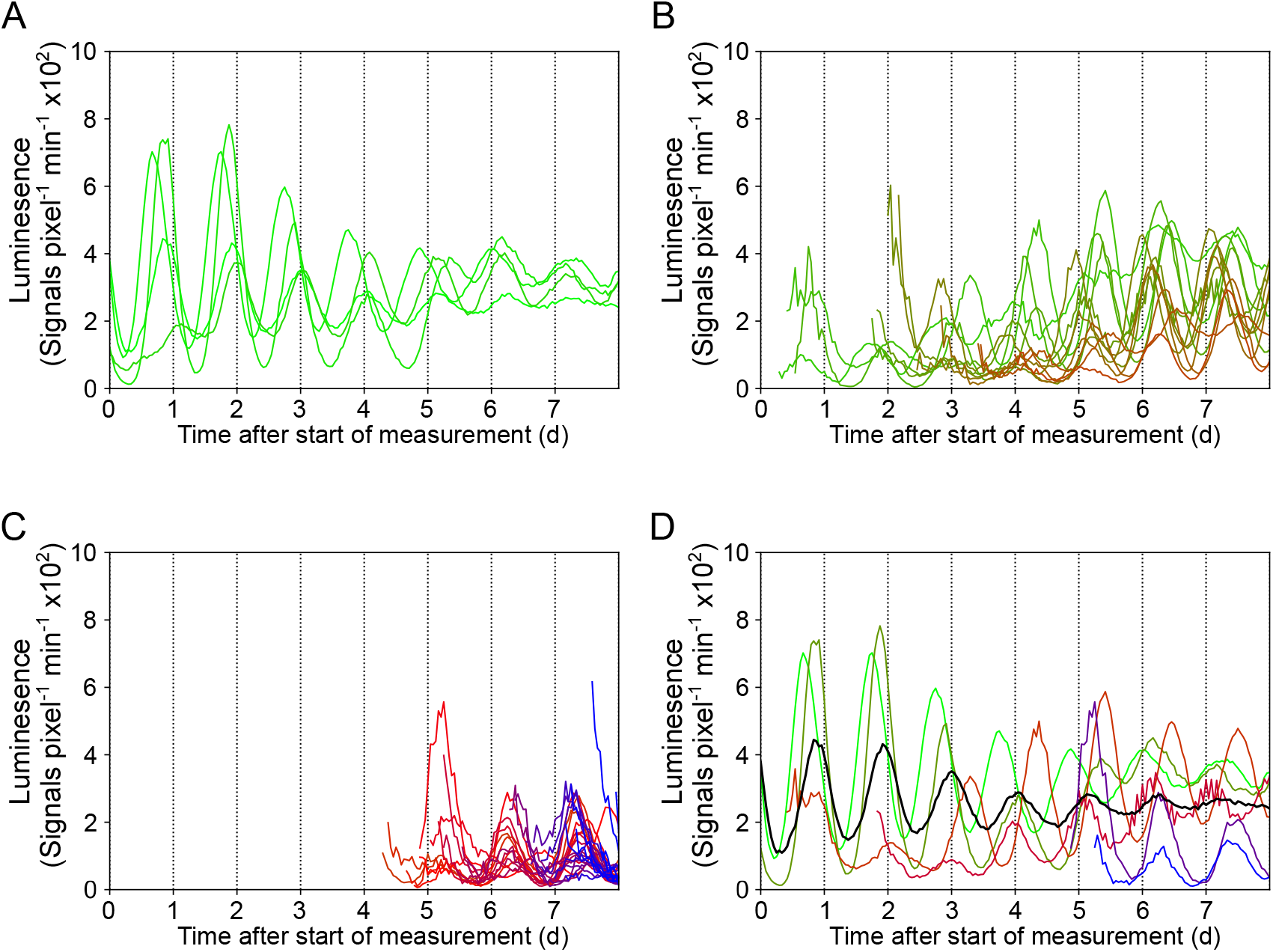
Bioluminescence rhythms of individual fronds of similar ages from an LL-grown plant in the experiment LLLL1. Bioluminescence rhythms plotted in Fig. 2A are classified into those of (A) fronds of the initial colony, (B) newly emerging fronds in the first 4 days, and (C) newly emerging fronds after 4 days. (D) Bioluminescence rhythms of a parental frond and its daughter fronds The black and colored lines represent the rhythm of the parent, LLLL1_1, and those of its daughters (1-1, 1-2, 1-3, 1-4, 1-5), respectively. Color-codes for (A-C) are the same as those in Fig. 2A. Colors in (D) are recoded.

**Fig. S7:**
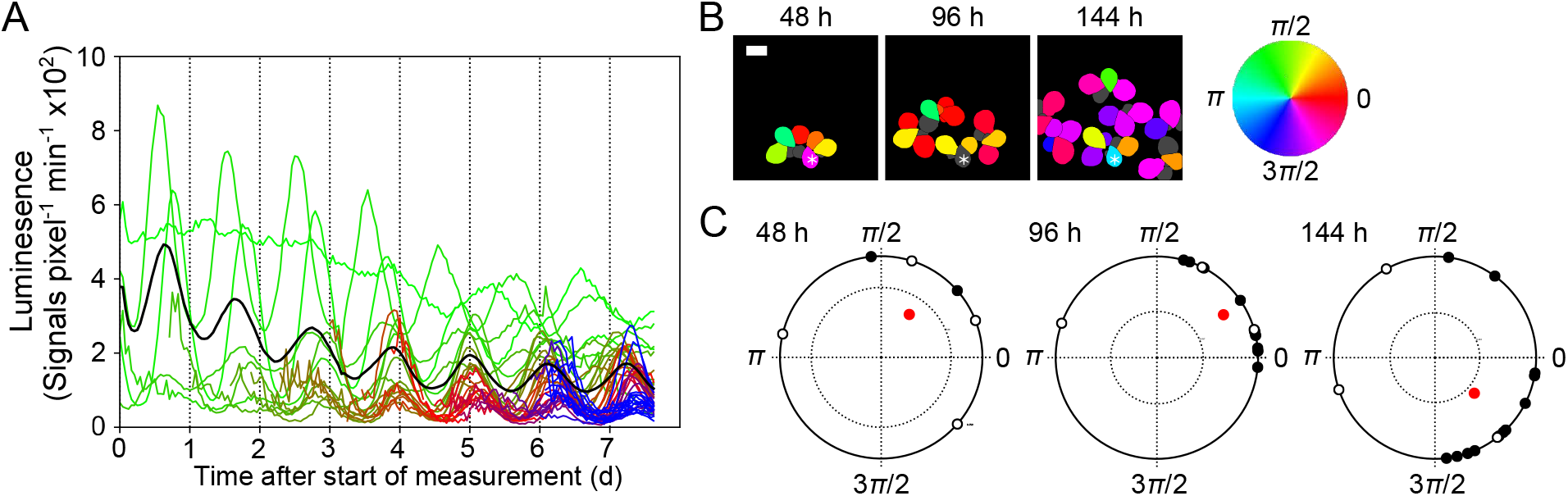
Bioluminescence rhythms and their phase properties of individual fronds from an LL-grown plant in the experiment LLLL2. Legends for (A), (B), and (C) are the same as those of Fig. S5, except for the definition of time 0: the start of measurement.

**Fig. S8:**
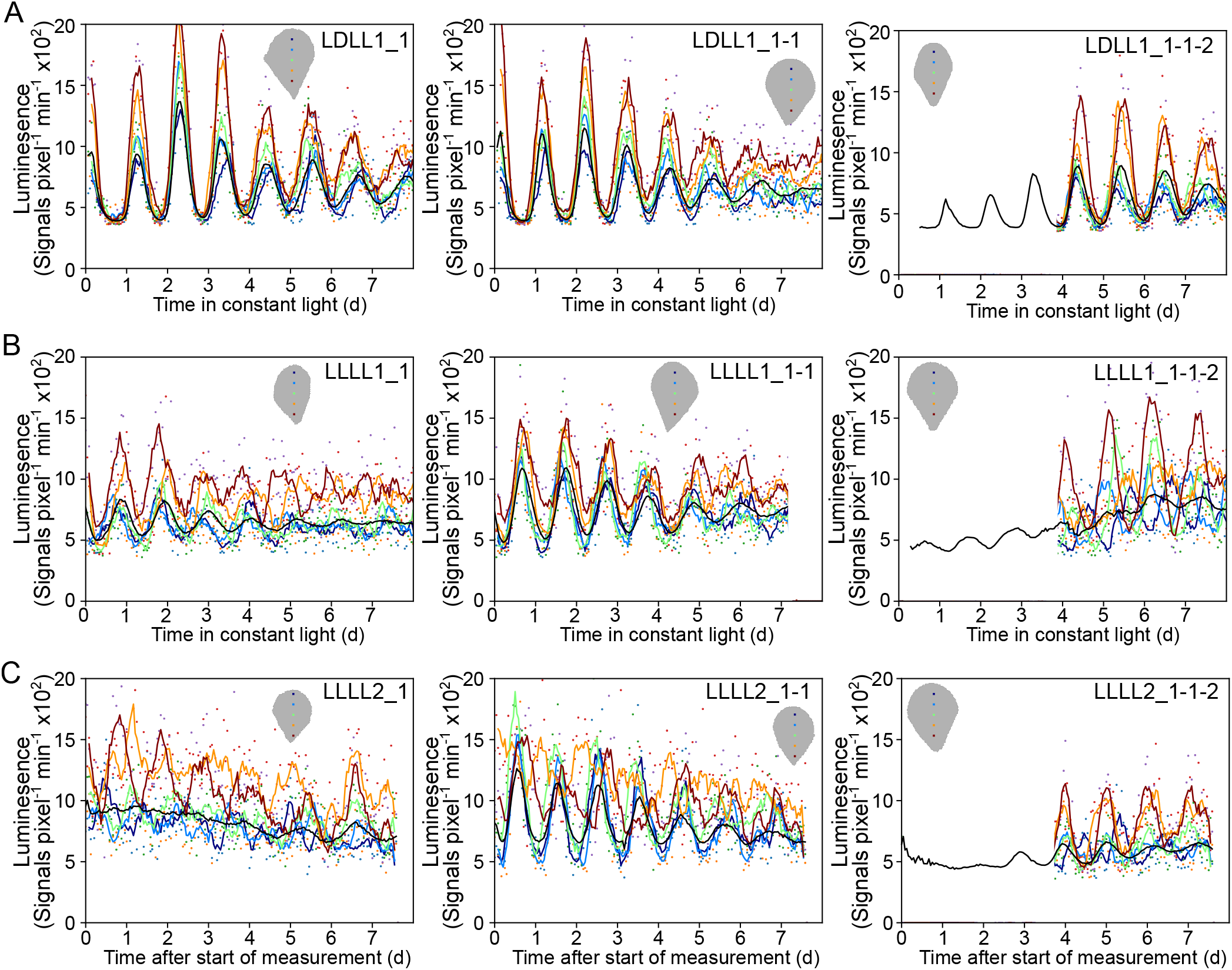
Luminescence rhythms of single pixels. Five pixels along the major axis of a frond are selected, and luminescence signals including background are plotted for three fronds of different ages in experiments (A) LDLL1, (B) LLLL1, and (C) LLLL2. In each graph area, a frond indicating the five pixels is drawn. These pixels locate every 15 pixels and include the geometric center. The lines represents the 3 h moving average for each pixel. Color gradation from red to blue represents the positions of pixels from the base (frond pocket side) to the top.

**Fig. S9:**
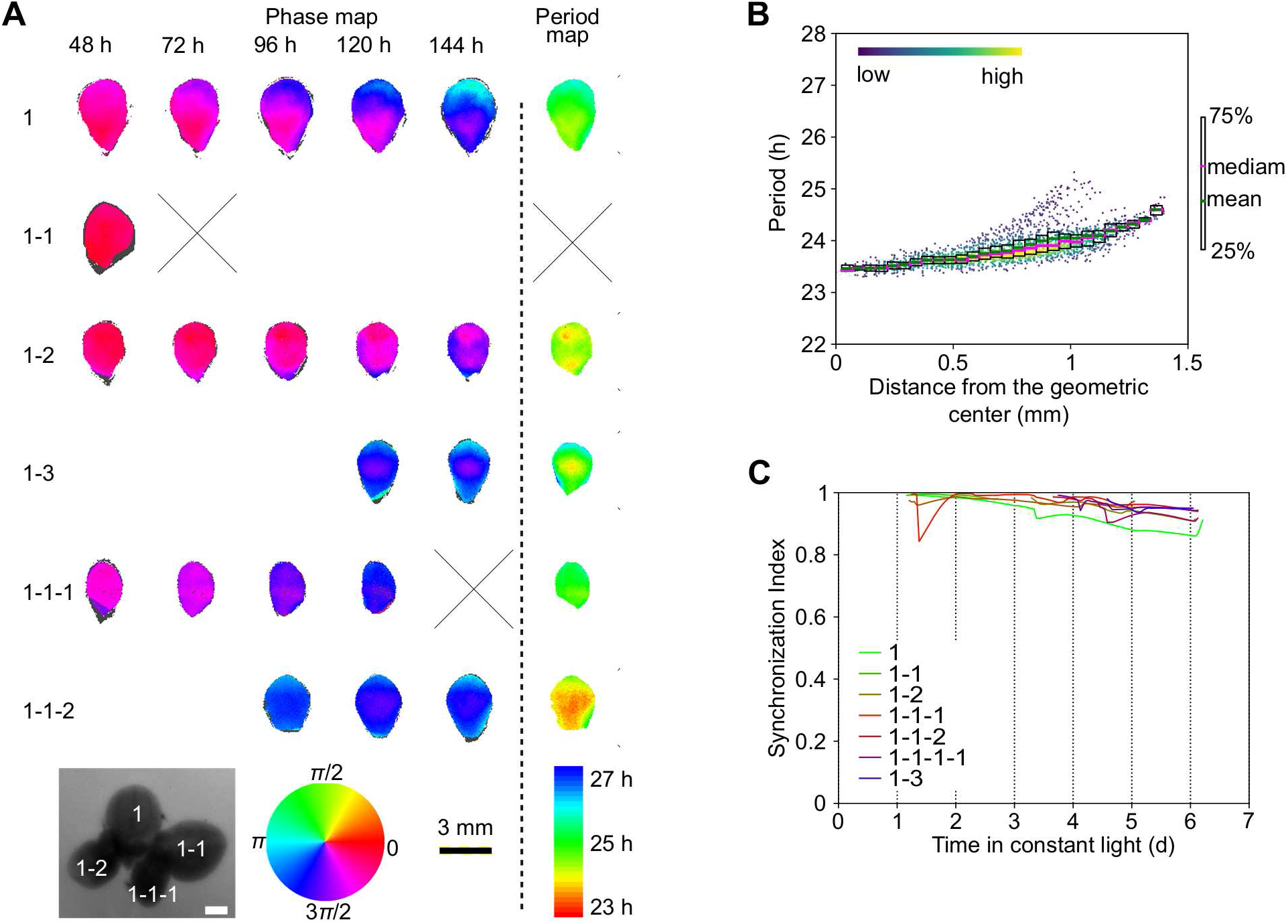
Characterization of bioluminescence rhythms in fronds from an LD-grown plant at the pixel level. The time series of luminescence of individual pixels in the experiment LDLL2 was used for analysis. (A) Phase maps and period maps. Phases at the indicated time and periods of circadian rhythms in each pixel of a frond are color-coded. Frond names are indicated on the left. A frond name has two parts: the number at the end and the remaining part. The number at the end represents the order of emergence from the parent whose name is represented in the remaining part. Top-view image of the starting plant with frond names is shown. The white bar is 1 mm. Period lengths were calculated using time series for 72 h starting from 24 h or the time at which the frond matured. Gray color represents pixels the phase or period of which was not defined. A cross mark represents the missing data due to the movement of the frond out of the visual field of the camera. (B) Relation between period lengths and distances from the geometric center of the frond (1-1-2). For each pixel, the period length and the distance from the geometric center to its position are plotted as dots. The density of dots is color-coded. Statistics (mean and quartiles) for the period lengths are plotted at every 0.05 mm interval of distances from the geometric center. (C) Temporal changes in SIs of individual fronds.

**Fig. S10:**
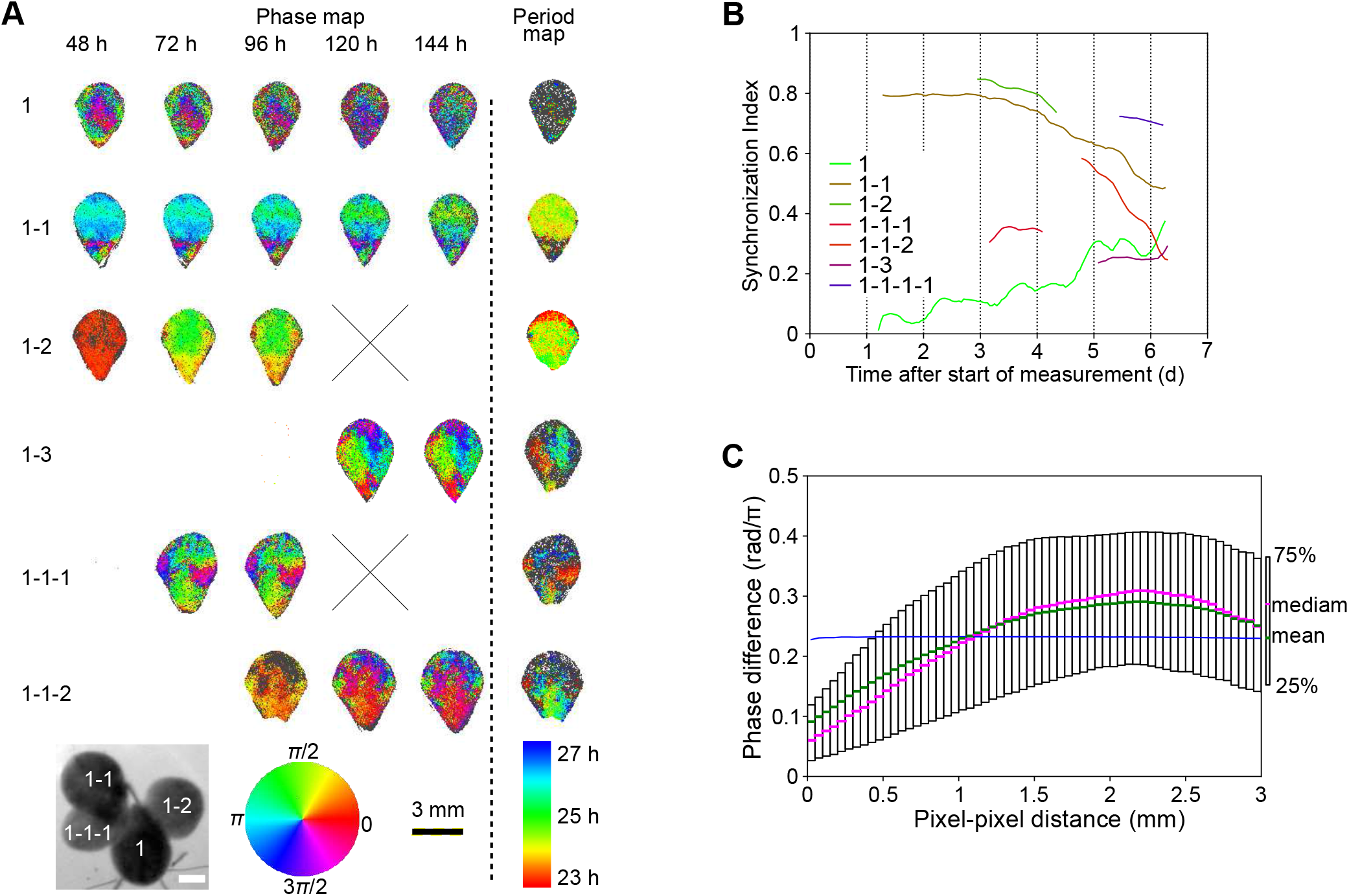
Characterization of bioluminescence rhythms in fronds from an LL-grown plant at the pixel level. The time series of luminescence of individual pixels in the experiment LLLL2 was used for analysis. (A) Phase maps and period maps. The legend is the same as that in Fig. S9A. (B) Temporal changes in SI of individual fronds. (C) Relationship between the phase differences between pixels and pixel-to-pixel distances. Phase differences for all pairs of pixels in the frond (1-3) at 144 h and pixel-to-pixel distances corresponding to the pairs were calculated. Statistics (mean and quartiles) for the phase differences are plotted at every 0.05 mm interval of pixel-to-pixel distances. Blue symbols at each pixel-to-pixel distance interval represent 95% confidence limits of the median, assuming samples in each interval have the same median value. The confidence limits were calculated based on resampling (10,000 repeats) for all pairs.

**Fig. S11:**
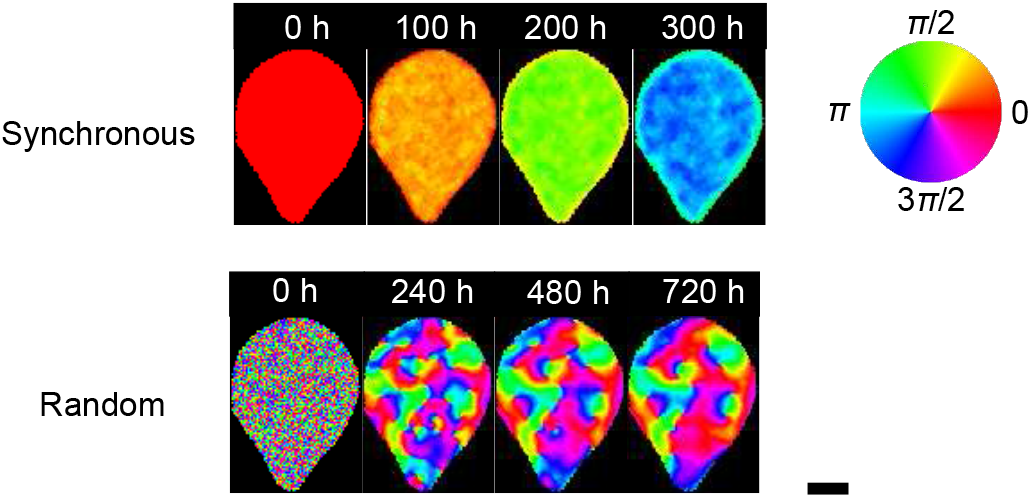
Time evolution of phase patterns in a frond via model simulation. Phase maps for a frond with synchronous- or random-phased oscillators at time 0 are drawn in the same simulation as in Fig. 4. Phases at the indicated time of circadian rhythms in each pixel of a frond are color-coded. The bar is 1 mm.

**Fig. S12:**
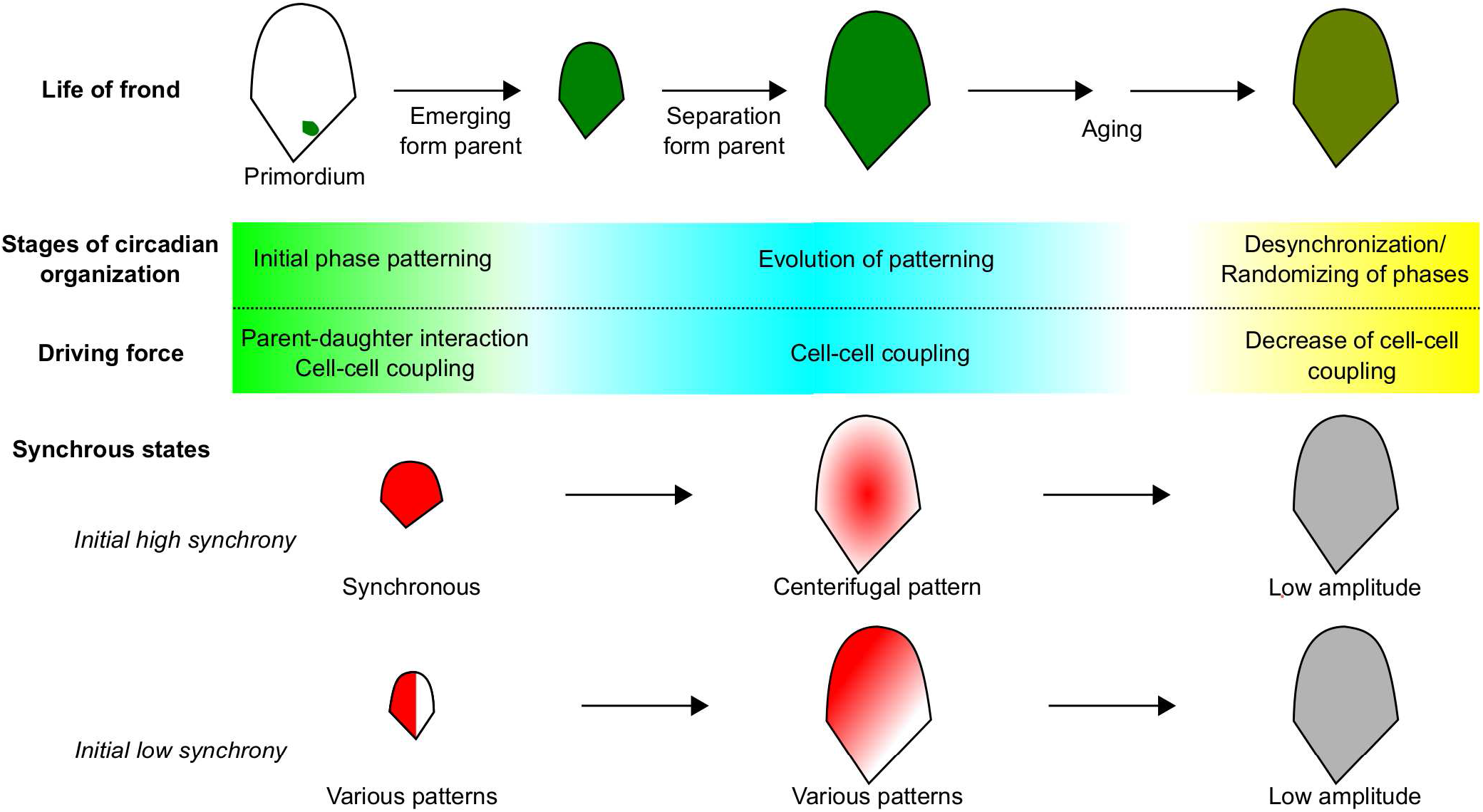
Hypothetical framework for spontaneous phase patterning in fronds of proliferating plants. The explanation is mentioned in the text.

**Table S1.**
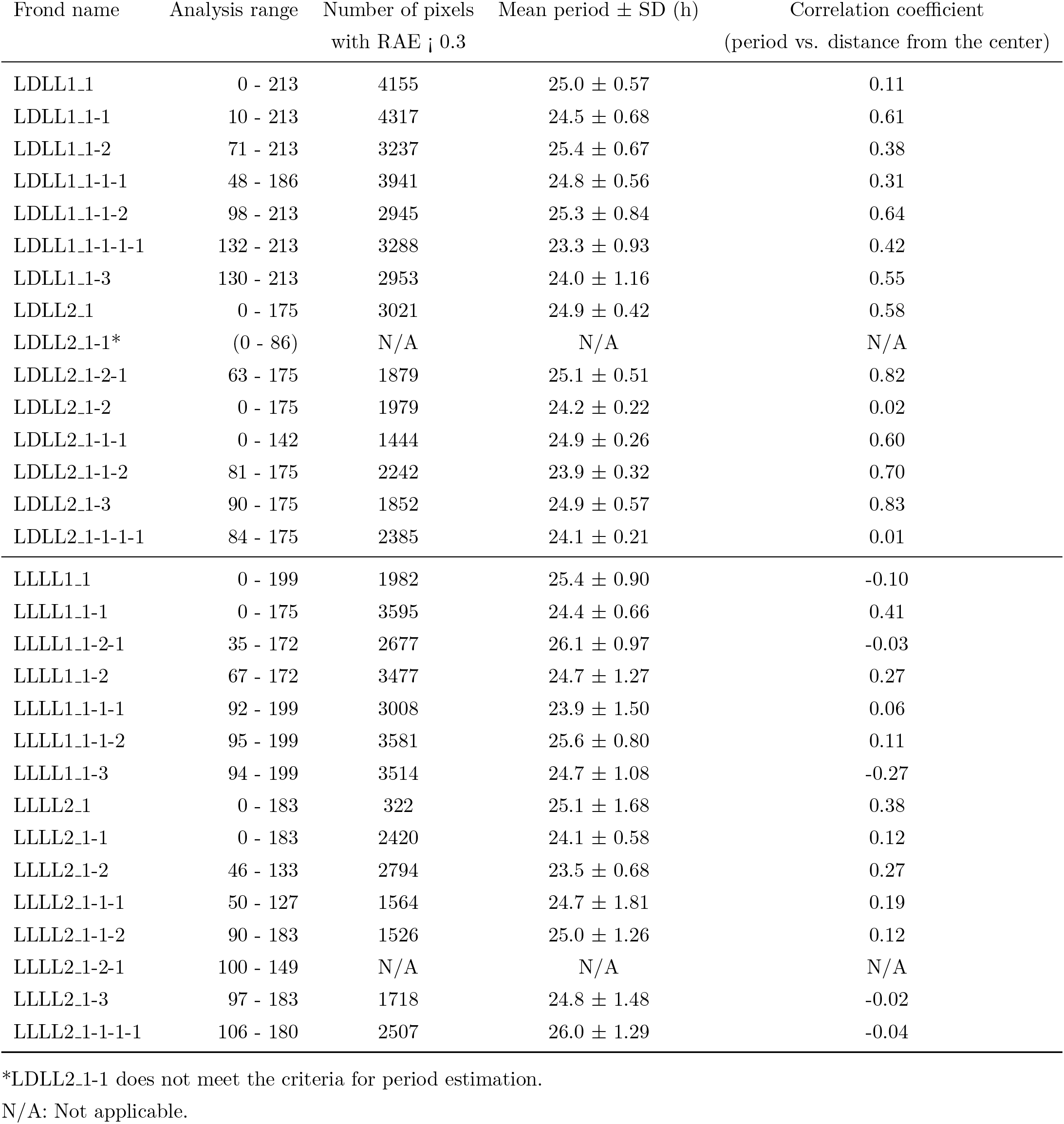
Periods of individual fronds at the pixel level.

Movie S1. Time-lapse bioluminescence imaging of LD-grown duckweed plants under constant light (Experiment LDLL1). (not uploaded)

Movie S2. Time-lapse bioluminescence imaging of LD-grown duckweed plants under constant light (Experiment LDLL2). (not uploaded)

Movie S3. Time-lapse bioluminescence imaging of LL-grown duckweed plants under constant light (Experiment LLLL1). (not uploaded)

Movie S4. Time-lapse bioluminescence imaging of LL-grown duckweed plants under constant light (Experiment LLLL2). (not uploaded)

Movie S5. Centrifugal phase pattern in the frond LLLL1_1-1. Phases of circadian rhythms in each pixel are color-coded. (not uploaded)

Movie S6. Transversal phase pattern in the frond LLLL1_1-3. Phases of circadian rhythms in each pixel are color-coded. (not uploaded)

Movie S7. Spiral phase pattern in the frond LLLL1_1-1-1. Phases of circadian rhythms in each pixel are color-coded. (not uploaded)

SI Dataset S1. Information of measurement for individual fronds of every experiment. (not uploaded)

## Notes

Author declaration: The authors declare no conflict of interest.

### Competing Interest Statement

The authors have declared no competing interest.

